# Safety, Immunogenicity and Efficacy of a Recombinant Vesicular Stomatitis Virus Vectored Vaccine Against Severe Fever with Thrombocytopenia Syndrome Virus and Heartland Bandaviruses

**DOI:** 10.1101/2021.11.29.470508

**Authors:** Philip Hicks, Jonna B. Westover, Tomaz B. Manzoni, Brianne Roper, Gabrielle L. Rock, Kirsten M. Boardman, Dallan J. Blotbter, Brian B. Gowen, Paul Bates

## Abstract

Severe fever with thrombocytopenia syndrome virus (SFTSV) is a recently emerged tickborne virus in east Asia with over 8,000 confirmed cases. With a high case fatality ratio, SFTSV has been designated a high priority pathogen by the WHO and the NIAID. Despite this, there are currently no approved therapies or vaccines to treat or prevent SFTS. Vesicular stomatitis virus (VSV) represents an FDA-approved vaccine platform that has been considered for numerous viruses due to its low sero-prevalence in humans, ease in genetic manipulation and promiscuity in incorporating foreign glycoproteins into its virions. In this study, we developed a recombinant VSV (rVSV) expressing the SFTSV glycoproteins Gn/Gc (rVSV-SFTSV) and assessed its safety, immunogenicity and efficacy in mice. We demonstrate that rVSV-SFTSV is safe when given to immunocompromised animals and is not neuropathogenic when injected intracranially into young immunocompetent mice. Immunization of *Ifnar*^-/-^ mice with rVSV-SFTSV resulted in high levels of neutralizing antibodies and protection against lethal SFTSV challenge. Additionally, passive transfer of sera from immunized *Ifnar*^-/-^ mice into naïve animals was protective when given pre- or post-exposure. Finally, we demonstrate that immunization with rVSV-SFTSV cross protects mice against challenge with the closely related Heartland virus despite low neutralizing titers to the virus. Taken together, these data suggest that rVSV-SFTSV is a promising vaccine candidate.

**Importance:** Tick borne diseases are a growing threat to human health. Severe fever with thrombocytopenia syndrome (SFTS) and Heartland viruses are recently recognized, highly-pathogenic, tick-transmitted viruses. The fatality rates for individuals infected with SFTSV or HRTV are high and there are no therapeutics or vaccines available. The recent introduction of the tick vector for SFTSV (*Haemaphysalis longicornis)* to the eastern half of the United States and Austrailia raises concerns for SFTSV outbreaks outside East Asia. Here we report the development of a potential vaccine for SFTSV and HRTV based on the viral vector platform that has been successfully used for an Ebola vaccine. We demonstrate that the rVSV-SFTSV protects from lethal SFTSV or HRTV challenge when given as a single dose. We evaluated possible pathogenic effects of the vaccine and show that it is safe in immune compromised animlas and when introduced into the central nervous system.

## Introduction

Severe fever with thrombocytopenia syndrome (SFTS) is an emerging tickborne disease caused by the SFTS virus (SFTSV, recently renamed *Dabie bandavirus*, formerly known as *Huaiyangshan Banyangvirus*). First identified in 2011 in China, SFTSV is a novel bunyavirus that can cause fever, thrombocytopenia, and leukocytopenia in infected individuals (1, 2). Subsequent reports later showed that SFTSV had been causing human disease since 2009 with a high case fatality ratio of approximately 30% (1, 2). Retrospective studies have found SFTSV to be endemic to, and causing disease in, South Korea, Japan, and Vietnam (3-5). Recent literature indicates the case fatality ratios range from 6-20% depending on the country studied, and disease progression is characterized by hemorrhagic tendency with fatal cases exhibiting multi-organ failure (6). In 2009 a novel related bunyavirus named Heartland virus (HRTV) was discovered in Missouri exhibiting a similar disease progression and transmission cycle to SFTSV (7).

The recently discovered SFTSV and HRTV are both bandaviruses in the order *Bunyavirales* and family *Phenuiviridae*. These viruses have a trisegmented, single-stranded RNA genome encoding 4 proteins. The S segment is ambisense and encodes the nucleoprotein (N) in the negative sense and a nonstructural protein (NSs) in positive sense (1). The L and M segments are negative sense and encode the RNA-dependent RNA polymerase (RdRp) and envelope glycoproteins, respectively. The glycoprotein is translated as a polyprotein which is proteolytically cleaved into two subunits, Gn/Gc (1, 8). The Gn/Gc complex recognizes its receptor or binding factor and entry is mediated by the fusion peptide within Gc (9-11). While studies have failed to isolate a receptor for SFTSV and HRTV Gn/Gc, some molecules have been identified as important binding or entry factors, including DC-SIGN and UGCG (9, 10, 12). Antibodies against both Gn and Gc have been shown to inhibit viral entry into cells (10, 13, 14).

The primary vector for SFTSV is the tick *Haemaphysalis longicornis* found throughout eastern Asia (15, 16). However, SFTSV has been found in other tick genera as well including *Ixodes* and *Amblyoma* (17, 18) suggesting that numerous tick species might transmit this pathogen. Due to transmission resulting from contact with ticks, SFTSV generally infects rural farmworkers. In recent years, the geographic distribution of *H. longicornis* has expanded and now includes Australia, New Zealand, and the United States, presenting further opportunities for SFTSV to spread (19-22). Although likely more rare, other transmission routes have been shown to be possible for SFTSV. Ferret studies have shown that SFTSV can be transmitted in the absence of ticks between co-housed ferrets or ferrets co-housed with a separator (23). The detection of SFTSV in ferret saliva, feces, and urine suggests that these fluids are a likely route of SFTSV transmission in the absence of ticks (23). Indeed, one report indicates that SFTSV can spread between humans in a nosocomial setting through contact with patient blood (24). Another case report showed likely zoonotic transmission to a human through the bite of an infected cat (25). Due to the high potential of SFTSV to cause deadly outbreaks, it has been categorized as a high priority pathogen for the development of vaccines and therapeutics by both the World Health Organization (WHO), and the National Institute of Allergy and Infectious Diseases (NIAID) (26, 27).

Currently no approved therapeutics or vaccines exist for use against SFTSV. This is in part due to lack of adequate animal models. Wild-type mice do not develop severe disease upon challenge thus alternatives must be used (28-30). Animals that succumb to SFTSV infection include cats, aged ferrets, *Stat2*^*-/-*^ hamsters, and *Ifnar*^*-/-*^ mice (28, 31-34). Despite these difficulties, several groups have designed and tested SFTSV vaccines, primarily using *Ifnar*^*-/-*^ mice. Tested vaccine platforms include DNA, virus-vectored, and attenuated recombinant SFTSV vaccines (35-39). These vaccines vary in their effectiveness and come with drawbacks. Here, we focus on developing and characterizing a recombinant vesicular stomatitis virus (rVSV) vaccine.

The livestock pathogen vesicular stomatitis virus (VSV) is generally non-pathogenic to humans and of low sero-prevalence (40, 41). Additionally, VSV is a powerful vaccine platform with genetically tractable models and a promiscuity to incorporate foreign glycoproteins in the virion (42). An often cited detriment of rVSV vaccines is the propensity for VSV to be neurotropic. It is, however, known that neuropathogenicity is conferred by the tropism of the viral glycoprotein (43, 44). Currently, the rVSV vaccine platform is approved for use against Ebola virus (EBOV) and has been successfully distributed in Africa during recent EBOV outbreaks (45, 46). Due to the proven nature of the rVSV platform, we made an rVSV-SFTSV virus containing the SFTSV Gn/Gc glycoprotein in place of the cognate VSV glycoprotein VSV-G.

It has been previously reported by another group that rVSV-SFTSV confers protective immunity to *Ifnar*^-/-^ mice (38). To go beyond what has been previously shown, we demonstrate that our rVSV-SFTSV is non-neurotropic and safe in immunocompromised animals. We also show that a single administration of vaccine virus is sufficient to induce protection against SFTSV challenge. Additionally, rVSV-SFTSV vaccintion induces high levels of antibodies in wild-type animals suggesting it can effectively be used in immune competent animals. Both therapeutic and prophylactic passive transfer of sera from immunized animals leads to protection upon challenge of unvaccinated animals suggesting antibodies correlate with protection against SFTSV. Finally, we demonstrate that our rVSV-SFTSV vaccine is cross-protective upon lethal HRTV challenge.

## Results

### rVSV-SFTSV is attenuated in *Ifnar*^*-/-*^ mice and exhibits a favorable safety profile

One safety concern with rVSV vaccines is the potential for neurotropism (47, 48). VSV-G, the cognate glycoprotein of VSV, is sufficient to catalyze viral entry into neurons, and VSV can replicate efficiently in neuron-like cells both *in vitro* and *in vivo* (49, 50). In addition, other rhabdoviruses such as rabies virus are known human pathogens that cause lethal neurotropic infections (51). It is currently unclear whether neurons can be infected by SFTSV or viruses harboring SFTSV glycoproteins. Neurologic symptoms have been observed in human SFTS cases, but it remains unclear whether the SFTSV glycoproteins can initiate entry into neurons *in vivo* (52). To test the neuropathogenic potential of rVSV-SFTSV, we injected escalating doses of rVSV-SFTSV or VSV Indiana strain intracranially into the right cerebral hemisphere of 4-week-old C57BL/6 mice and observed the mice for 14 days. All groups of mice lost 2-5% body weight the day following intracranial injection independent of inoculum composition (Fig. 1A). Weight loss in mice injected with rVSV-SFTSV was reversed by 4 days post-infection (dpi). In contrast, weight loss was more severe in mice challenged with VSV and survivors exhibited slower recovery. None of the mice injected with rVSV-SFTSV met humane endpoints during the experiment (Fig. 1B). In contrast, lethal disease was seen in mice injected with VSV that trended with inoculum dose. Neurologic effects were quantified by using a neurologic sign scoring scale that ranged from 0 (no neurologic signs) to 4 (severe neurologic signs). No neurologic signs were observed in mice inoculated with rVSV-SFTSV or vehicle. In contrast, a range of neurologic manifestations were observed in most mice injected with VSV (Fig. 1C). All the observed neurologic effects occurred between 2-7 dpi (data not shown).

**Figure 1.**
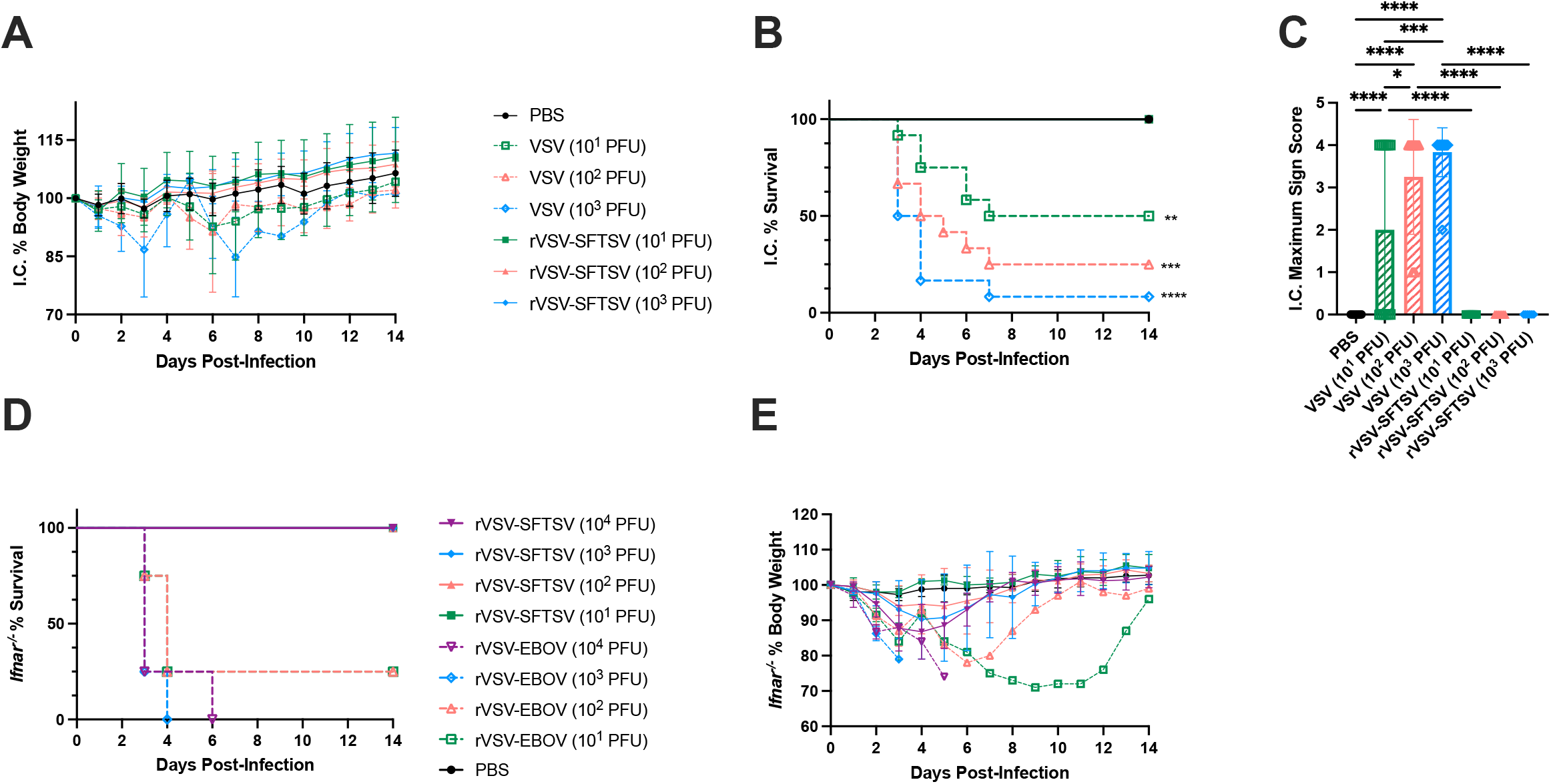
rVSV-SFTSV has a favorable safety profile compared to rVSV-EBOV and parental VSV. (A) Weight change, (B) survival proportions, (C) and maximal neurologic disease severity score in C57BL/6 mice challenged intracranially (IC) with 10^1^, 10^2^, or 10^3^ PFU of parental VSV or rVSV-SFTSV into the right cerebral hemisphere (Mantel-Cox test and ordinary one-way ANOVA; *, P<0.0332; **, P<0.0021; ***, P<0.0002; ****, P<0.0001) (D) Survival proportions and (E) weight loss of *Ifnar*^*-/-*^ mice challenged intraperitoneally with PBS or 10^1^, 10^2^, 10^3^, or 10^4^ PFU of either rVSV-SFTSV or rVSV-EBOV. Weight changes were reported as percentages of body weight measured immediately pre-challenge. (Mantel-Cox test; *, P<0.0332; **, P<0.0021; ***, P<0.0002; ****, P<0.0001).

The rVSV-EBOV vaccine received FDA approval despite severe pathogenicity in *Ifnar*^*-/-*^ and *Stat1*^*-/-*^ immunocompromised mice (47, 48). To evaluate the pathogenicity of rVSV-SFTSV, we challenged groups of *Ifnar*^*-/-*^ mice with escalating doses of rVSV-SFTSV or rVSV-EBOV. All mice injected with rVSV-SFTSV were alive 14 dpi (Fig. 1D). In contrast, at least 50% of mice infected with rVSV-EBOV met humane endpoints by 4 dpi, and all mice challenged with at least 10^3^ PFU succumbed by 6 dpi. All groups of mice challenged with rVSV-EBOV exhibited weight loss beginning 2 dpi which progressed with time (Fig. 1E). The surviving mice challenged with rVSV-EBOV lost at least 20% body weight prior to their recovery. Mice challenged with rVSV-SFTSV also began losing weight by 2 dpi, but the weight loss was less severe compared to rVSV-EBOV groups. Of the groups of mice challenged with rVSV-SFTSV, only the group infected with 10^4^ PFU lost more than 10% body weight. Recovery from weight loss began between 4-5 dpi for all mice challenged with rVSV-SFTSV. Collectively, these experiments showed that rVSV-SFTSV is not neuropathogenic and demonstrated a significantly more favorable safety profile than the currently approved rVSV-EBOV vaccine in immunocompromised mice.

### Single vaccination with rVSV-SFTSV induces high levels of neutralizing antibodies

To functionally characterize humoral responses to rVSV-SFTSV vaccination we assessed antibody neutralization potential by focus reduction neutralization titer of 50% (FRNT_50_) in several animal models. To assess whether the lethal *Ifnar*^*-/-*^ mouse model could mount an antibody response, mice were immunized intraperitoneally (IP) with 10^2^, 10^3^, or 10^4^ plaque forming units (PFU) of rVSV-SFTSV. At 21 dpi sera were collected and analyzed for FRNT_50_. Approximately half of the animals vaccinated with 10^2^ PFU failed to seroconvert (Fig. 2A). Increasing the vaccination dose increased rates of seroconversion and neutralization titers (Fig. 2A). These titers are promising given thay previous work on influenza and SARS-CoV-2 suggest that neutralizing titers of 40-80 are sufficient for protection (53, 54).

**Figure 2.**
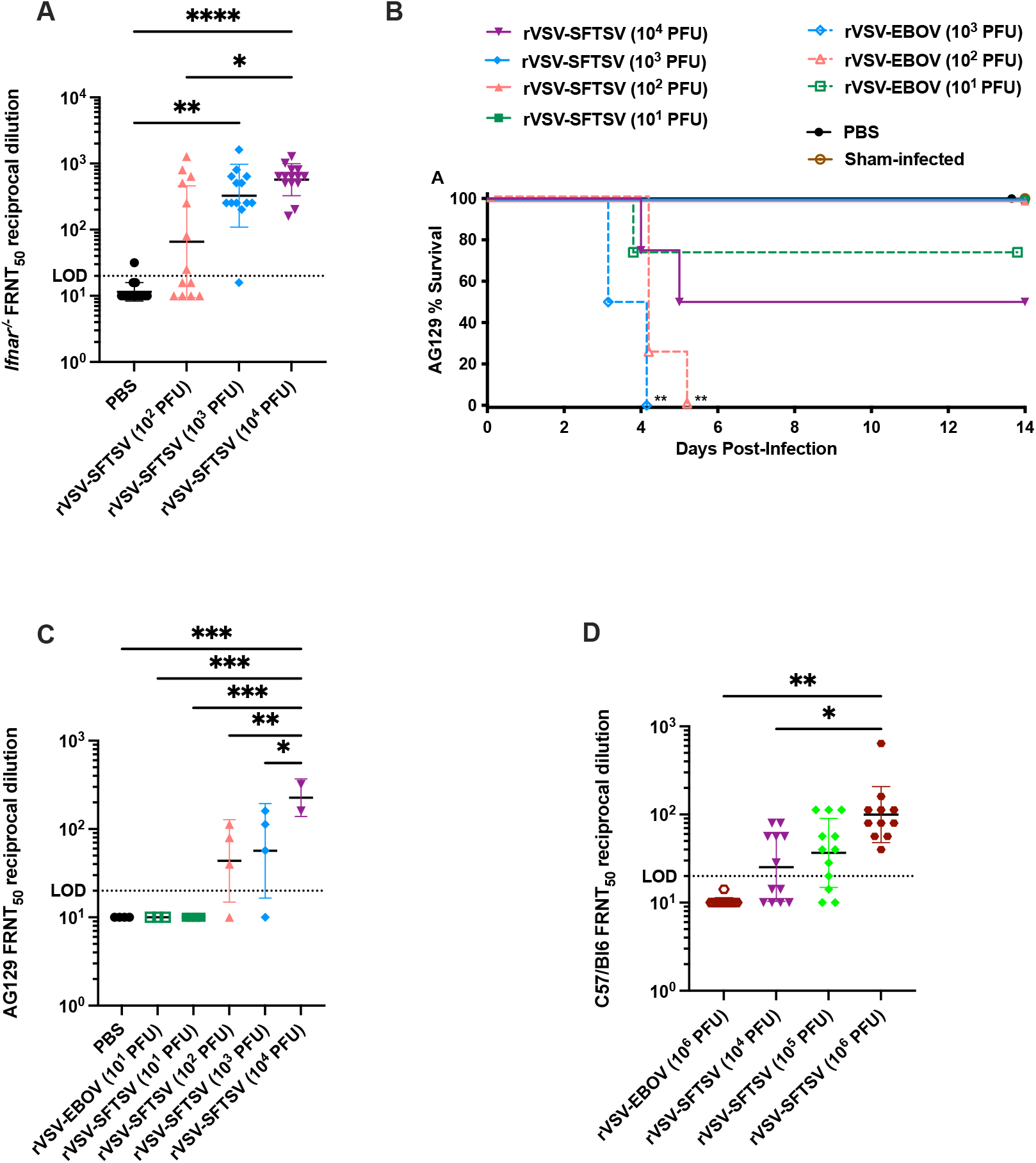
rVSV-SFTSV induces neutralizing antibodies across different mouse strains. (A) *Ifnar*^*-/-*^ mice were immunized with PBS, 10^2^, 10^3^, or 10^4^ PFU rVSV-SFTSV. Serum neutralizing antibodies were quantified by measuring 50% decrease in pseudovirus foci, the reciprocal endpoint dilution is shown (Ordinary one-way ANOVA; *, P<0.0332; **, P<0.0021; ***, P<0.0002; ****, P<0.0001). (B,C) AG129 mice were vaccinated with varying doses of rVSV-SFTSV and monitored for survival (B) and had serum collected 21 days post vaccination and FRNT_50_ was assessed (C) (Mantel-Cox test and ordinary one-way ANOVA; *, P<0.0332; **, P<0.0021; ***, P<0.0002; ****, P<0.0001). (D) Wild-type C57/Bl6 mice were immunized with rVSV-SFTSV and had serum neutralization titers determined at 21 days post treatment (Ordinary one-way ANOVA; *, P<0.0332; **, P<0.0021; ***, P<0.0002; ****, P<0.0001).

The high levels of neutralizing antibodies achieved with vaccination of *Ifnar*^*-/-*^ mice was somewhat surprising as interferons (IFN)s are important drivers of immune responses. To determine whether mice deficient in both type I and type II IFN receptors also elicit high levels of neutralizing antibodies, we immunized AG129 (IFN-α/β and γ receptor-deficient) mice with 10^1^-10^4^ PFU of rVSV-SFTSV. Notably, 2 of 4 mice immunized with 10^4^ PFU rVSV-SFTSV succumbed to viral infection (Fig. 2B). Mice receiving 10 PFU rVSV-SFTSV failed to generate a neutralizing antibody response (Fig. 2C). Animals receiving higher doses had mean neutralizing titers ranging from 60 to 240 with increasing dosage (Fig. 2C). These results demonstrate that rVSV-SFTSV elicits humoral responses even in highly immunocompromised animals lacking type I and II IFN responses.

It is well documented that VSV infection is highly sensitive to IFN responses (55, 56). To determine whether rVSV-SFTSV can induce a neutralizing antibody response in immunocompetent mice, we immunized C57BL/6 mice with 10^4^, 10^5^, or 10^6^ PFU of rVSV-SFTSV. Dosages were increased relative to *Ifnar*^*-/-*^ mice to account for IFN responses interfering with rVSV-SFTSV replication and thus reducing the humoral immune response in the immune competent mice. In contrast to what was seen with the immune deficient mice, no weight loss was observed in the C57BL/6 mice at any vaccine dose (data not shown). Additionally, despite the increased dosages, neutralizing titers were far lower than those observed in *Ifnar*^*-/-*^ mice suggesting that rVSV-SFTSV is sensitive to IFN, consistent with previous reports (Fig. 2D). A dosage dependent increase in FRNT_50_ titers was observed with mice receiving 10^6^ PFU rVSV-SFTSV achieving a mean titer of 113 (Fig. 2D). Notably, all mice immunized with 10^6^ PFU rVSV-SFTSV sero-converted (Fig. 2D). These data indicate that despite VSV’s sensitivity to IFN, rVSV-SFTSV induces responses in immunocompetent animals that reach neutralizing titers predicted to be protective.

### rVSV-SFTSV protects *Ifnar*^*-/-*^ mice from lethal SFTSV challenge and reduces viral titers in tissues

Because of the high neutralizing antibody titers measured in *Ifnar*^*-/-*^ mice vaccinated with rVSV-SFTSV, we hypothesized that the vaccine would protect these mice against lethal SFTSV challenge. To test this hypothesis, vaccinated mice were challenged subcutaneously with 10 PFU of SFTSV 23 days post-vaccination (2 days after blood collection for neutralizing antibody titration). A single group of unvaccinated mice received 8 days of 100 mg/kg/day of favipiravir therapy following SFTSV challenge as a positive control for protection (57). As expected, all mice vaccinated with PBS succumbed by 8 dpi (Fig. 3A). In contrast, only 60% of mice vaccinated with 10^2^ PFU, and all *Ifnar*^*-/-*^ mice vaccinated with at least 10^3^ PFU, survived the lethal SFTSV challenge. Mice vaccinated with at least 10^3^ PFU were protected from weight loss following SFTSV challenge while rapid weight loss was observed in PBS-vaccinated mice beginning by 3 dpi (Fig. 3B). Mild weight loss occurred post-challenge in mice that received 10^2^ PFU of vaccine, but this trend was driven primarily by the three individuals that succumbed to disease. Vaccination-associated weight loss was also observed in this experiment and was consistent in magnitude to that shown previously (Fig. 1E).

**Figure 3.**
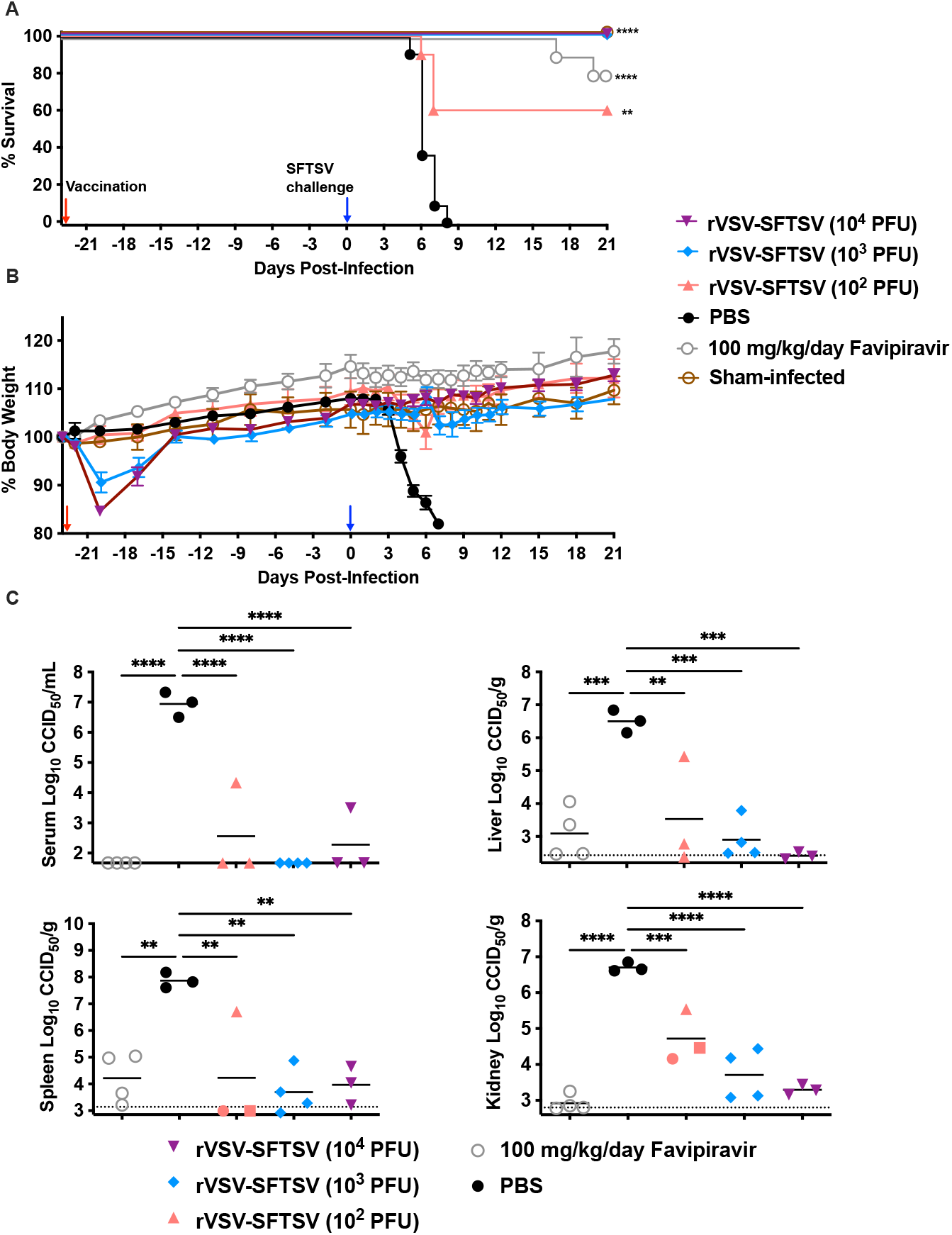
Vaccination with rVSV-SFTSV protects *Ifnar*^*-/-*^ mice from lethal SFTSV challenge. (A) Survival proportions and (B) percent weight change in *Ifnar*^*-/-*^ mice challenged subcutaneously with 10 PFU SFTSV (blue arrow) 23 days after intraperitoneal vaccination with PBS, or 10^2^, 10^3^, or 10^4^ PFU rVSV-SFTSV (red arrow). Weight change is reported as percentage change in body weight relative to starting weight prior to vaccination. One group of mice received favipiravir daily for eight days following SFTSV challenge to serve as a positive control for protection. (Mantel-Cox test; *, P<0.0332; **, P<0.0021; ***, P<0.0002; ****, P<0.0001). (C) SFTSV titers in serum liver, spleen, and kidney of mice five days post-challenge in mice subjected to the same vaccination schedule as those in (A) and (B). Horizontal dotted lines indicate the limit of detection of the assay (Ordinary one-way ANOVA; *, P<0.0332; **, P<0.0021; ***, P<0.0002; ****, P<0.0001).

To assess the effect rVSV-SFTSV vaccination has on SFTSV viremia and tissue viral loads, groups of 4 mice were vaccinated and challenged in, parallel following the timeline described above. These subsets of mice were sacrificed 5 days following SFTSV challenge and serum, liver, spleen, and kidney were collected for SFTSV quantification by endpoint titration on Vero E6 cells. All groups of vaccinated mice had significantly reduced SFTSV serum and tissue viral titers (Fig. 3C). In the liver and kidney, there was a trend towards dose-dependence with mice vaccinated with the highest dose of rVSV-SFTSV having the lowest viral titers. Favipiravir treatment also reduced SFTSV titers compared to mice vaccinated with PBS. These data demonstrate that rVSV-SFTSV does not provide sterilizing immunity to SFTSV challenge but rather reduces replication in the vaccinated animals.

### Passive serum transfer imparts protective immunity to naïve mice

Although our data demonstrate that rVSV-SFTSV vaccination induces neutralizing antibody response and also protects against lethal SFTSV infection, it remains unclear if antibody-mediated immunity is sufficient to confer SFTSV protection. To assess whether antibodies alone impart protection against lethal SFTSV infection, we performed passive serum transfer. 60 µl or 20 µl of sera from rVSV-SFTSV-immunized or negative control *Ifnar*^*-/-*^ mice were administered to naïve *Ifnar*^*-/-*^ mice either prophylactically (2 days prior to challenge) or therapeutically (2 days post challenge). The pooled sera used for passive transfer had an approximate FRNT_50_ titer of 450, while the FRNT_50_ titer for negative control sera was below the limit of detection. Approximately 33% of *Ifnar*^*-/-*^ mice receiving 60 µl of immune sera prophylactically were protected against lethal SFTSV challenge (Fig. 4A). When 60 µl of immune sera was administered therapeutically, 62% of mice survived (Fig. 4A). Animals given 20 µl of immune sera were not protected from challenge regardless of when sera were administered (Fig. 4A). All mice in the positive control group receiving 10^3^ PFU rVSV-SFTSV 7 days prior to challenge survived (Fig. 4A).

**Figure 4.**
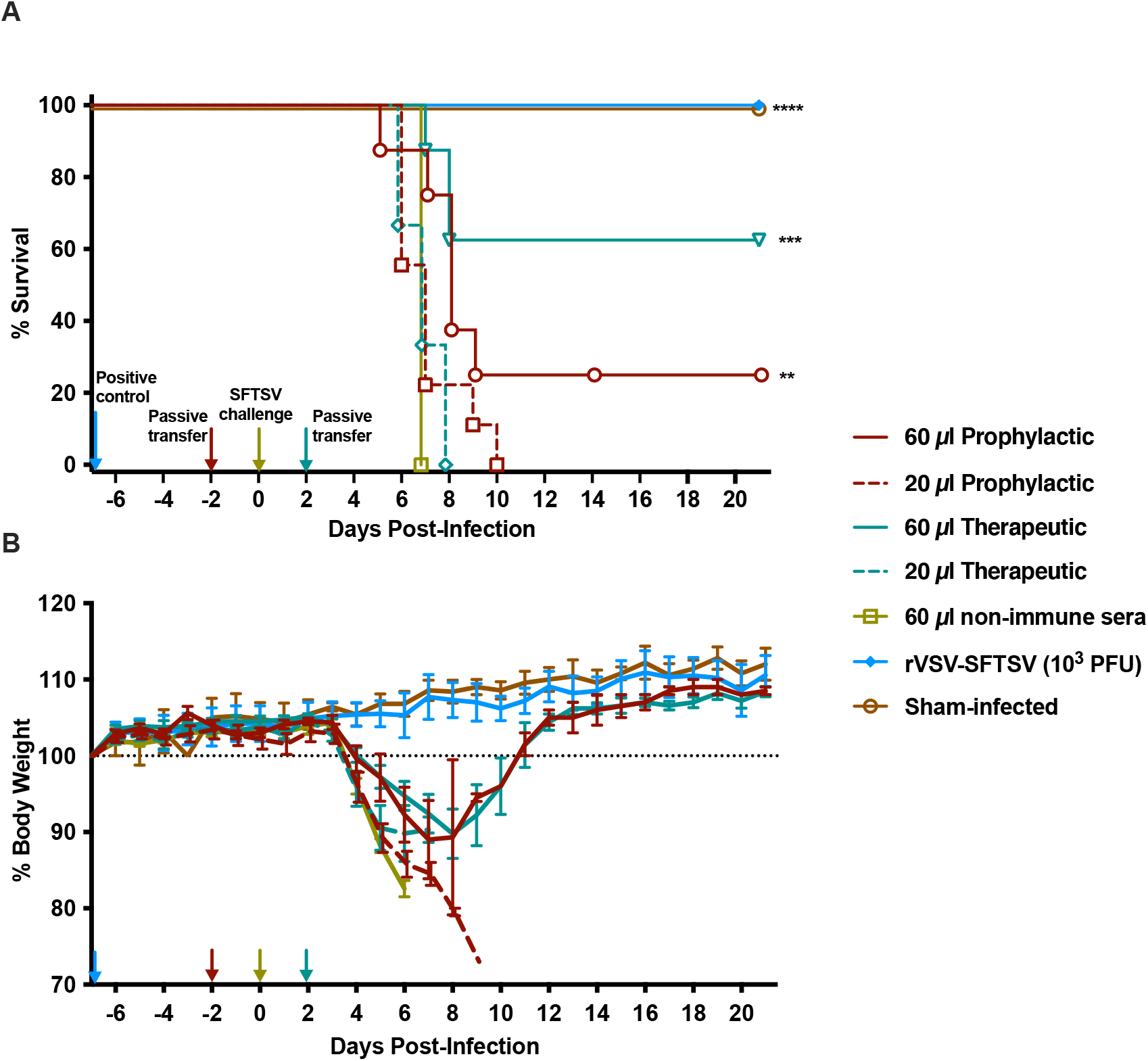
Passive transfer of sera from immunized mice protects naïve mice against SFTSV challenge. Survival (A) and weight loss (B) curves are shown from naïve animals receiving immune sera either 2 days prior to or 2 days post challenge with 10 PFU of SFTSV. Mice immunized with 10^3^ PFU of the rVSV-SFTSV 7 days prior to challenge served as the positive control. Blue arrow, immunization with rVSV-SFTSV 7 days prior to challenge; Red arrow, passive transfer 2 days prior to challenge; Yellow arrow, SFTSV challenge; Teal arrow, passive transfer 2 days post SFTSV challenge. (Mantel-Cox test; *, P<0.0332; **, P<0.0021; ***, P<0.0002; ****, P<0.0001).

Animal weights measured daily during the study positively correlated with the survival data (Fig. 4B). The most dramatic weight loss after the SFTSV challenge occurred in the group treated with the non-immune sera and the groups treated with the lower quantities of immune sera. All surviving mice treated with the 60 µl dose of immune sera recovered fully from the infection after losing approximately 10% body weight (Fig. 4B). The mice vaccinated with 10^3^ PFU rVSV-SFTSV did not experience any weight loss due to the vaccine virus or upon SFTSV infection (Fig. 4B).

### rVSV-SFTSV vaccination cross protects against lethal HRTV challenge

Next, we evaluated whether vaccination with rVSV-SFTSV confers cross protection against HRTV challenge. In AG129 mice, a dose of 10^4^ PFU rVSV-SFTSV was partially lethal (Fig. 2B). Thus, we modified immunization dosages to 10^2^, 10^2.5^, and 10^3^ PFU rVSV-SFTSV. Mice were then challenged with a lethal dose of a mouse-adapted HRTV (MA-HRTV) 21 days post vaccination. A group of unvaccinated mice were treated with 100 mg/kg/day favipiravir for 8 days following MA-HRTV challenge. Eighty percent of the mice that received the two highest doses of 10^2.5^ or 10^3^ PFU rVSV-SFTSV survived the challenge, with 60% of the mice vaccinated with the lowest dose (10^2^ PFU) also surviving (Fig. 5A). As expected, all PBS-vaccinated mice succumbed to MA-HRTV disease by 8 dpi and all the favipiravir-treated animals were protected (Fig. 5A). Most of the infected mice experienced considerable weight loss beginning 4 to 6 days post MA-HRTV challenge (Fig. 5B). Surviving mice fully recovered and had weight gain comparable to favipiravir-treated animals.

**Figure 5.**
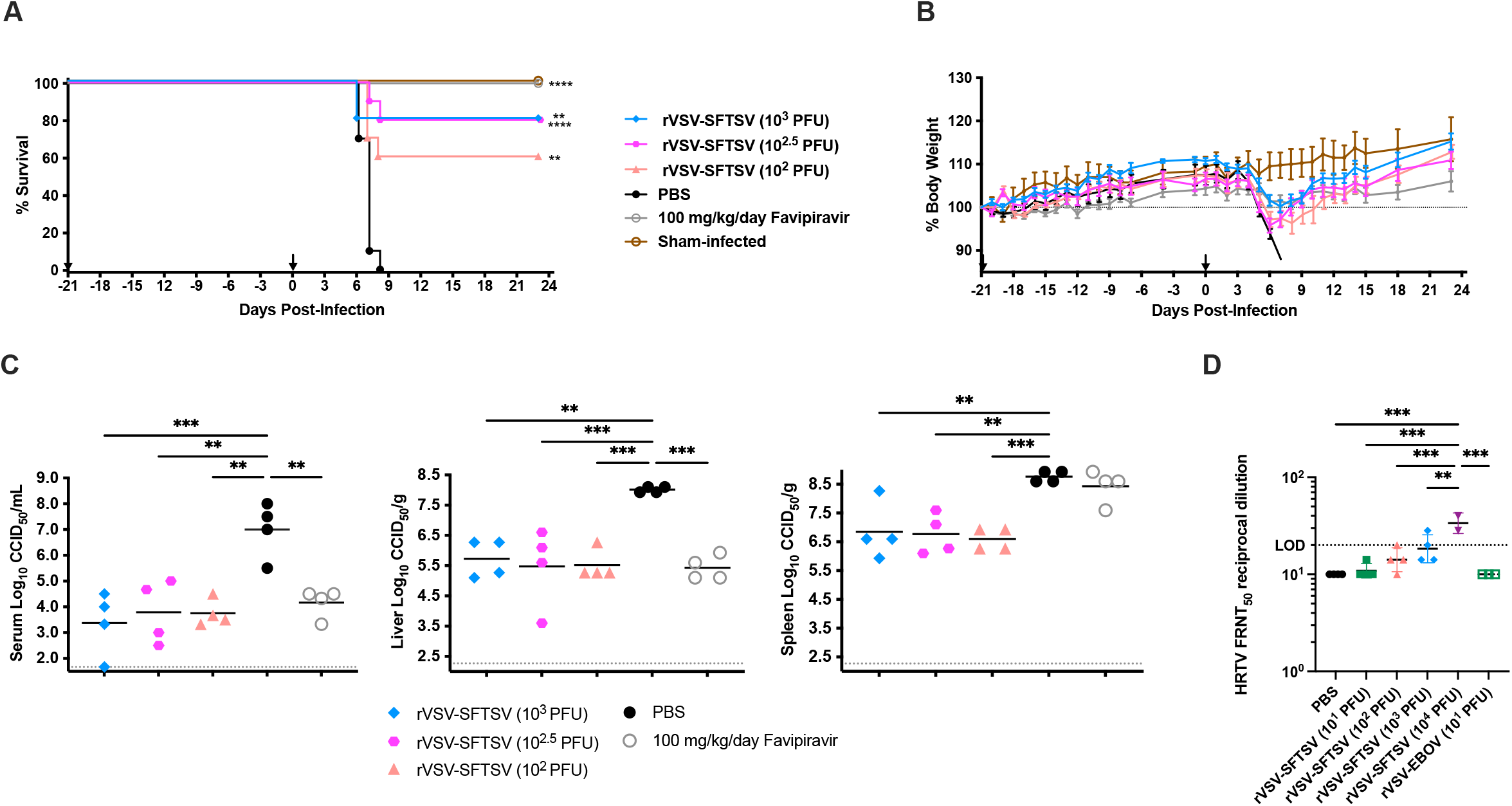
rVSV-SFTSV vaccination cross-protects animals against MA-HRTV challenge. AG129 mice were IP immunized with escalating doses of rVSV-SFTSV then challenged with MA-HRTV 21 days post immunization. (A) Survival and (B) weight loss curves are shown from immunization until completion of experiment. Black arrows indicate vaccination and challenge times at −21 and 0 days respectively (Mantel-Cox test; *, P<0.0332; **, P<0.0021; ***, P<0.0002; ****, P<0.0001). (C) Four animals in each vaccination group were sacrificed 5 days post challenge to assess serum, liver, and spleen virus titers (Ordinary one-way ANOVA; *, P<0.0332; **, P<0.0021; ***, P<0.0002; ****, P<0.0001) (D) Sera was collected from subsets of animals 21 days post immunization and prior to HRTV challenge. Sera was analyzed for neutralizing antibodies against HRTV using a pseudotyped virus with the HRTV Gn/Gc glycoprotein (Ordinary one-way ANOVA; *, P<0.0332; **, P<0.0021; ***, P<0.0002; ****, P<0.0001).

Subsets of 4 mice per experimental group were sacrificed on day 5 post MA-HRTV challenge for collection of blood, liver, and spleen tissue for measurement of viral loads by endpoint titration using an infectious cell culture assay. Mice immunized with rVSV-SFTSV had significantly reduced MA-HRTV titers comparable to the favipiravir positive control (Fig. 5C). In parallel, serum from vaccinated animals was analyzed for FRNT_50_ to determine whether cross neutralization of HRTV occurred. Mice vaccinated with 10^2^ or 10^3^ PFU rVSV-SFTSV had moderate SFTSV neutralization titers of 40 and 60, respectively, with one mouse in each group failing to seroconvert (Fig. 2C). Mice receiving 10^4^ PFU immunization doses reached higher neutralization titers against SFTSV but 2 of 4 animals succumbed the vaccine virus (Fig. 2B, C). Interestingly, cross neutralization of HRTV was only observed in sera from 2 of 4 mice receiving 10^3^ PFU immunizations and the surviving 10^4^ PFU immunized animals from the safety and immunogenicity study (Fig. 5D). In mice immunized with 10^3^ PFU rVSV-SFTSV, neutralizing titers against HRTV were at or just above the limit of detection indicating weak cross reactivity (Fig. 5D). Sera from mice receiving lower immunization doses did not have neutralization activity despite partial protection from MA-HRTV challenge (Fig. 5A, C, D). Lack of cross neutralization titers suggests that the protective effect in the context of survival and reduced viral loads may be due to cell-mediated immunity or other non-neutralizing antibody functions.

## Discussion

SFTSV is an emerging public health threat in southeast Asia with case fatality rates ranging from 4 to 30%. This high variability in case fatality rates may reflect access to health care, the genetic background of infected populations, and the virulence of the infecting SFTSV strain (58-60). Given that no therapeutics or vaccines are available to curtail or prevent an outbreak of SFTS and that the virus is transmitted by multiple tick species with expanding geographic ranges, the threat SFTSV poses to public health is significant (18, 61). This has caused the WHO to list SFTSV in its prioritized pathogen research blueprint and the NIAID to include it as a category C priority pathogen (26, 27). In response to the potential threat from SFTSV, many SFTSV vaccines are being developed using a variety of different technologies including protein subunit, DNA, and recombinant viral platforms (35, 36, 38, 39, 62, 63).

*Ifnar* ^*-/-*^ mice are susceptible to lethal SFTSV challenge and have therefore been used to evaluate an array of SFTSV vaccines that use DNA, protein subunit, and recombinant viral technologies (36, 38, 39, 62). A DNA vaccine encoding the ectodomains of Gn and Gc and a chimeric N-NSs fusion protein on a single plasmid only protected 40% of mice from lethal challenge after three doses of vaccine (62). However, a similar vector that contained an additional open reading frame encoding IL-12 was fully protective using an identical vaccination schedule (62). Interestingly, neither regimen induced neutralizing antibodies. This same group also developed a protein subunit vaccine by fusing Gn and Gc to the Fc region of human immunoglobulin heavy chain to create Gn-Fc and Gc-Fc, respectively (62). Vaccination of mice with either Gn-Fc or Gc-Fc in alum induced neutralizing antibody titers greater than 1:100 in a FRNT_50_ assay. This vaccination regimen, however, achieved only 50% and 0% protection from lethal SFTSV challenge with Gn-Fc or Gc-Fc, respectively (62). Recombinant vaccinia virus and VSV technologies have also been explored. LC16m8 strain recombinant vaccinia virus vaccines encoding SFTSV N alone, SFTSV Gn/Gc alone, or both N and Gn/Gc in combination fully protected against lethal SFTSV challenge (36). These vaccines however failed to induce FRNT_50_ neutralizing titers greater than 1:40, and not surprisingly, the vaccine encoding N alone failed to induce detectable neutralizing activity. Of the vaccine platforms evaluated thus far in *Ifnar* ^*-/-*^ mice, recombinant VSVs induced the highest neutralizing titers. In our study, mean FRNT_50_ neutralizing titers ranging from 282 to 642 were induced by a single dose of rVSV-SFTSV HB29 strain that ranged from 10^2^ - 10^4^ PFU. A similar rVSV encoding the SFTSV AH12 strain glycoprotein sequence elicited a neutralizing titer of 682 following vaccination with a single dose of 2 × 10^4^ PFU (38).

Aged ferrets have also been used as a lethal SFTSV model to study the efficacy of both DNA vaccines and live-attenuated vaccines harboring mutations in NSs (35, 39). Representative vaccines from these platforms that fully protected ferrets from lethal challenge included a combination of two plasmids that encoded SFTSV Gn and Gc, a combination of five plasmids that encode SFTSV Gn, Gc, RdRp, N, and NSs, a recombinant SFTSV HB29 strain virus encoding the P102A substitution in NSs, and a recombinant SFTSV HB29 strain virus encoding an NSs containing a truncated carboxy-terminus of 12 amino acids. Both DNA vaccine platforms are given in a three dose regimen and elicit modest levels of neutralizing antibodies. Passive transfers from animals vaccinated against SFTSV Gn/Gc were protective suggesting neutralizing antibodies are a correlate of protection (35). It remains unclear what contribution cellular responses make to protection from SFTSV challenge. Unfortunately, the age-related changes that render ferrets susceptible to lethal SFTSV challenge are unknown. This makes it difficult to directly compare results obtained from aged ferrets to other SFTSV animal models.

The only currently FDA-approved rVSV vaccine, rVSV-EBOV, is highly pathogenic and lethal in *Ifnar* ^*-/-*^ mice. In contrast, rVSV-SFTSV only caused mild-to-moderate weight loss at doses that elicited protective immunity. Unlike the parental VSV vector, rVSV-SFTSV did not cause neurologic disease when injected intracranially into 4-week-old C57BL/6 mice, suggesting that this vaccine strain is not neurotropic. Despite these promising results, it is possible that rVSV-SFTSV may be too attenuated in animals containing a functional IFN system. This possibility is supported by the lower neutralizing antibody titers measured in vaccinated C57BL/6 mice compared to those in *Ifnar* ^*-/-*^ mice. However, neutralizing serum levels in vaccinated C57BL/6 mice were higher than titers that are typically expected to protect against other viruses such as influenza (53). This uncertainty highlights the need for immune-competent animal models to evaluate the efficacy of SFTSV vaccines and therapeutics.

Neutralizing antibodies elicited in the context of a functional humoral response are thought to be a correlate of protection against lethal SFTS in humans (64). As such, passive transfer of sera from vaccinated animals has been previously shown to be protective when used prophylactically in an *Ifnar* ^*-/-*^ mouse model of lethal SFTSV (38). Extending these findings, our results demonstrated that serum from mice vaccinated with rVSV-SFTSV could protect against lethal SFTSV challenge when used either prophylactically or therapeutically suggesting the protective capacity of Gn/Gc-targeted antibodies against SFTSV. In support of the role for antibodies in protection against SFTSV disease, a monoclonal antibody recognizing Gn protects *Ifnar*^*-/-*^ mice from lethal challenge (65). The efficacy of neutralizing sera administered therapeutically has implications for monoclonal antibody or antibody cocktail management of patients with SFTS.

In addition to direct neutralization by elicited antibodies, our data also suggest other mechanisms for protection against SFTSV pathogenesis. In our passive transfer studies, following redistribution of transferred sera from the peritoneal cavity to the circulation and subsequent dilution within the host’s blood, it is likely that the neutralizing titer of the recipient’s sera would be below the limit of detection of our FRNT_50_ assay. Still, the dose of antibodies contained within 60 μL of donor sera was sufficient to protect mice under both prophylactic and therapeutic conditions. Surprisingly, we observed better protection when sera were transferred therapeutically (2 dpi) compared to prophylactically (−2 dpi). While the mechanisms underlying this observation are unknown, it is possible that uncharacterized variables such as the pharmacokinetics and bioavailability of the sera components responsible for protection at sites of virus replication may be responsible. In addition, our results suggest that passive transfer intervention may be beneficial in areas endemic for SFTSV within high-risk populations for severe SFTS that have reported recent tick bites but are yet to show symptoms. Further studies using appropriate animal models should be conducted to better evaluate the temporal relationship of passive transfer therapies with both infection (by experimental inoculation or tick bite) and clinical disease onset.

Protection against lethal challenge in the absence of high neutralizing antibody titers has been observed in SFTSV vaccines using recombinant vaccinia virus and DNA technologies (36, 62). These observations suggest that other mechanisms in addition to neutralizing antibodies may also contribute to protection against SFTS. In this study, we demonstrated that vaccination with rVSV-SFTSV either 7 or 21 days prior to SFTSV challenge was fully protective (Figures 3A and 4A). Protection at 7 days, a time at which humoral responses are not fully developed, suggests that mechanisms other than antibodies can be protective. Since the T cell immune response peaks at approximately 7 days post-vaccination, it is possible that these cells also contribute to the protection elicited by the vaccine (66, 67). Additionally, we noted protection against SFTSV lethality in some of the rVSV immunized mice where the neutralizing titer was below the limit of detection. rVSVs are known to stimulate robust T_H_1 immune responses in vaccinated animals and humans (68-70). The data presented here suggest that rVSV-SFTSV may also confer protection by inducing type 1 immunity and CD8+ T cells. It is also possible that NK cells or other components of the innate immune system are important mediators of this early protective effect of the rVSV vaccine. More studies will be required to elucidate the potential role of T cells following vaccination with rVSV-SFTSV or infection by authentic SFTSV.

HRTV is an emerging bandavirus closely related to SFTSV that can cause lethal disease (71, 72). Previous studies have shown that sera raised by vaccination against SFTSV glycoproteins can cross-neutralize viruses harboring HRTV glycoproteins (38). In addition, this same study showed that a rVSV-HRTV protects *Ifnar* ^*-/-*^ mice against lethal SFTSV infection. The present study is the first to report that rVSV-SFTSV protects *Ifnar* ^*-/-*^ mice against a lethal HRTV challenge. Despite protection against lethal disease, sera from rVSV-SFTSV-vaccinated mice neutralized viruses pseudotyped with HRTV glycoproteins much less efficiently than viruses pseudotyped with SFTSV glycoproteins. Of the four serum samples collected from AG129 mice vaccinated with 10^2^ PFU of rVSV-SFTSV, only one had a detectable neutralizing antibody titer (reciprocal dilution of 20) against VSV pseudotyped with HRTV glycoproteins. Despite this, 60% of AG129 mice that received this dose of vaccine survived MA-HRTV challenge. These data suggest that other systems stimulated by rVSV-SFTSV vaccination, such as a T_H_1 response, could have contributed to the protection against HRTV. Additionally, alternative effector functions of antibodies beyond neutralization may contribute to the protective effect described here. Further studies are needed to elucidate which immune system components are responsible for cross-protection as the results of these studies could inform future vaccine development for these recently emerged bandaviruses.

## Acknowledgements

We thank Ashley Dagley, Julio Landinez, Brittney Downs and Parker Webber for technical support with vaccine and passive transfer efficacy studies. This work was supported by the National Institutes of Health (R21AI142638, R01AI152236, T32AI055400, T32AI007324 and HHSN272201700041I).

## Author Contributions

P.B. and B.B.G. conceived and supervised the study. P.H., J.B.W., T.B.M., B.R., G.L.R., K.M.B. and D.J.B. performed the experiments. P.B., B.B.G., P.H., J.B.W., T.B.M., analyzed the data. P.H., J.B.W., T.B.M wrote the manuscript.

## Materials and Methods

### Cells, Viruses, and Mice

ATCC verified and mycoplasma free 293T and Vero E6 cells were maintained in DMEM (Corning, #10-013-CV) containing 10% FBS (Corning, #35-010-CV), and 2mM L-glutamine (Corning, #25-005-Cl). Cells were passaged every 2-3 days.

Recombinant viruses harboring an additional open reading frame encoding EGFP (refered to as VSV throughout this paper) in genomic position 5, or encoding heterologous viral glycoproteins in genomic position 4 (rVSV-SFTSV and rVSV-EBOV) were launched and described previously (12, 73). rVSV-SFTSV and rVSV-EBOV also contain an additional open reading frame in position five encoding mCherry. All recombinant viruses were grown in Vero E6 cells by infecting a confluent T-175 flask at an MOI of 0.3-0.5. Virus was collected at 48-72 hours post infection with the addition of Hepes buffer pH7.4 to 25mM. Media was clarified by centrifuging at 6000 times gravity for 5 minutes at 4 °C twice. Virus was then frozen at −80 °C until used for ultracentrifugation. Virus was concentrated by ultracentrifuging virus-containing media through a 20% sucrose gradient at 26,000rpm for 2 hours at 4 °C using SW-32 tubes in a Beckman Coulter Optima XPN-80 ultracentrifuge. After removal of the sucrose and media, pelleted virus was placed on ice with 500 µl hepes buffered saline overnight. The next day virus pellets were resuspended and frozen at −80 °C. Viral titer was determined by plaque assays on Vero E6 cells with a 1.25% Avicel RC-591 NF (DuPont, #RC591-NFBA500) overlay and then stained with 1% crystal violet.

SFTSV, strain HB29, was obtained from Dr. Robert Tesh (WRCEVA; World reference Center for Emerging Viruses and Arboviruses at the University of Texas Medical Branch, Galveston, TX). The virus stock (5.6 × 10^6^ PFU/ml; 1 passage in Vero E6 cells) used was from a clarified cell culture lysate preparation. Virus stock was diluted in sterile minimal essential medium (MEM) and inoculated by subcutaneous injection of 0.2 ml containing approximately 10 PFU.

The mouse-adapted HRTV (MA-HRTV) strain used was derived from the MO-4 strain obtained from Dr. Robert Tesh (WRCEVA). The MA-HRTV stock (4.7 × 10^8^ 50% cell culture infectious dose (CCID_50_/ml); 1 passage in Vero E6 cells, 5 passages in AG129 mice) used was prepared from clarified liver homogenate. The virus stock was diluted in sterile MEM and inoculated bilaterally in two IP injections of 0.1 ml each for a total inoculation of 40 CCID_50_.

C57BL/6 mice were ordered from Jackson Labs (Bar Harbor, ME). AG129 and *Ifnar*^*-/-*^ mice were obtained from breeding colonies at Utah State University. 4 week old C57BL/6 mice were used for intracranial challenge experiments. 8 week old C57BL/6 mice or *Ifnar*^*-/-*^ mice on the C57BL/6 background were used for all other experiments. All mouse experiments were done using equal numbers of male and female mice. All mice were given approximately 7 days to acclimate to their cages and vivarium prior to each experiment. Mice were weighed immediately prior to all vaccination and infection procedures. All mice were anesthetized using 1% isoflurane in air delivered by vaporizer (Northern Vaporisers, Skipton, UK) to the anesthesia chamber. Injection sites were first prepared by cleaning with a 70% ethanol pad. Intracranial injection experiments and some vaccination experiments without authentic SFTSV challenge were performed under animal biosafety level (ABSL) 2 conditions at the University of Pennsylvania. All other vaccination experiments that included authentic SFTSV challenged were performed in BSL3 conditions at Utah State University.

All animals were treated ethically using complying with guidelines set by the USDA and Utah State University Institutional Animal Care and Use Committee and the University of Pennsylvania Laboratory Animal Resources guidelines.

### Immunization

Vaccines were diluted to the desired concentrations with sterile PBS just prior to vaccination by IP injection. All immunizations were done with a 200 µl inoculum. Favipiravir, the positive control for the rVSV-SFTSV vaccine efficacy study, was kindly provided by the Toyama Chemical Co., Ltd. (Toyama, Japan) and prepared in a meglumine solution for administration by IP injection.

### Intracranial infection and neurologic sign scoring

To evaluate neuropathogenesis, 4-week-old C57BL/6 mice were injected intracranially into the right cerebral hemisphere using a 1ml Hamilton syringe with Repeating Syringe Dispenser (Hamilton Company, Reno, NV). Inocula contained 0, 10^1^, 10^2^, or 10^3^ PFU of rVSV-SFTSV or rVSV-EGFP and were diluted to a total injection volume of 10 μl with PBS. Mice we monitored during anesthesia recovery until they were ambulatory. Mice were weighed daily and were observed for neurologic signs. Neurologic signs were assigned a severity score ranging from 0-4. Mice scored “0” showed no signs of illness and were bright, alert, and responsive when handled. Mice scored a “1” showed mild signs of illness without clear signs of neurologic illness including body hunching, depressed activity, or mild grimace. Mice assigned a “1” had normal ambulation and responded normally to being handled. Mice assigned a “2” had clinical signs consistent with mild encephalitis including hyperexcitability or altered gait that did not impair linear locomotion and used all four limbs. Mice assigned a “3” had more severe neurologic signs which included paraparesis of one or two limbs, mild head tilt, and altered gait that did impair linear locomotion (such as spinning). Mice assigned a “4” had severe neurologic signs that were inconsistent with life including complete pelvic limb paraplegia, ataxia, or tremors/seizures. Mice scored with a “4” were humanely euthanized with CO_2_.

### Blood collection

Mice were isofluorane anesthetized and blood was collected through the submandibular route using Goldenrod lancets 5mm (Medipoint, Mineola, NY). Blood was maintained on ice after collection. Serum was separated from blood by centrifugation at 8,000 RPM for 30 minutes at 4 °C in an Eppendorf 5424R centrifuge (Eppendorf, Enfield, CT). Serum was heat inactivated by incubating at 56 °C for 30 minutes. While running neutralization assays, serum was stored at 4 °C, for long term storage serum was frozen at −80 °C.

### Pseudovirus neutralization assay

#### Production of VSV pseudotype with SFTSV Gn/Gc

293T cells plated 24 hours previously at 2 × 10^7^ cells per T-175 flask were transfected using Lipofectamine 2000 (Invitrogen, #11668-019) using manufacturers protocol. Briefly, tubes each containing 1.75ml optimem (Gibco, #31985-070) were made. In one tube 100ul of Lipofectamine 2000 reagent was added and gently mixed. In the other, 45ug of pCAG-SFTSV Gn/Gc expression plasmid was added, tubes were allowed to sit for 5 minutes at room temperature. Lipofectamine and DNA containing tubes of optimum were combined and gently mixed, after 20 minutes incubating at room temperature. Solution was added to flask of 293T cells, after 4 hours cells were fed with fresh media. Thirty hours after transfection, the SFTSV Gn/Gc expressing cells were infected for 2-4 hours with VSV-G pseudotyped VSVΔG-mNeon at an MOI of ∼1-3 (Generated by deleting the cognate VSV-G and linking mNeon to the n-terminus of P. Virus was launched as previously described (12)). After infection, the cells were washed twice with FBS-free media to remove unbound virus. Media containing the VSVΔG-mNeon SFTSV Gn/Gc pseudotypes was harvested 30 hours after infection and clarified by centrifugation twice at 6000g then aliquoted and stored at −80 °C until used for antibody neutralization analysis.

#### Antibody neutralization assay using VSVΔG-mNeon SFTSV Gn/Gc

Vero E6 cells were seeded in 100 μl at 2×10^4^ cells/well in a 96 well collagen coated plate. The next day, 2-fold serially diluted serum samples were mixed with VSVΔG-mNeon SFTSV Gn/Gc pseudotype virus (100-200 focus forming units/well) and incubated for 1hr at 37 °C. Also included in this mixture to neutralize any potential VSV-G carryover virus was 1E9F9, a mouse anti-VSV Indiana G, at a concentration of 600 ng/ml. The antibody-virus mixture was then used to replace the media on VeroE6 cells. 16 hours post infection, the cells were washed and fixed with 4% paraformaldehyde before visualization on an S6 FluoroSpot Analyzer (CTL, Shaker Heights OH). Individual infected foci were enumerated and the values compared to control wells without serum. The focus reduction neutralization titer 50% (FRNT_50_) was measured as the greatest serum dilution at which focus count was reduced by at least 50% relative to control cells that were infected with pseudotype virus in the absence of mouse serum. FRNT_50_ titers for each sample were measured in two to three technical replicates performed on separate days.

### Virus titer determination

Virus titers were assayed using an infectious cell culture assay as previously described (74). Briefly, a specific volume of tissue homogenate or serum was serially diluted and added to triplicate wells of Vero E6 (African green monkey kidney) cell monolayers in 96-well microtiter plates. The viral cytopathic effect (CPE) was determined 11 days after plating and the 50% endpoints calculated as described (75). The lower limits of detection were 1.67 log_10_ CCID_50_/ml serum and 2.43-3.14 log_10_ CCID_50_/g tissue. In samples presenting with virus below the limits of detection, a value representative of the limit of detection was assigned for statistical analysis.

### Passive transfer

The immune sera from mice vaccinated with rVSV-SFTSV (approximate FRNT_50_ of 453), non-immune sera, and recombinant vaccine rVSV-SFTSV (7.12 × 10^7^ PFU/ml) were diluted with sterile PBS so that the volume of each treatment was 100 µl. Sera was delivered by IP injection 2 days prior to or post challenge with SFTSV. Mice receiving the rVSV-SFTSV vaccine were immunized 7 days prior to challenge. Monitoring of mouse weight began at 7 days prior to challenge and continued 21 days post SFTSV challenge.

### Statistical and Data Analysis

The Mantel-Cox log-rank test was used for analysis of Kaplan-Meier survival curves. A one-way analysis of variance (ANOVA) with the Dunnett’s post test to correct for multiple comparisons was used to assess differences in virus titers. A one-way ANOVA with Tukey’s multiple comparisons post-hoc test was used to assess FRNT_50_ titers and maximum neurologic sign scores. All statistical evaluations were done using Prism 9 (GraphPad Software, La Jolla, CA).

